# Cellular Senescence Signatures Accurately Predict Clinical Phenotypes But Lack Uniqueness. Part I: Demonstrating Equivalence Classes of Senescence

**DOI:** 10.64898/2026.11.17.767150

**Authors:** S. Ma, M. Xu, L. Wang, E. Kummerfeld, M. Vilas, H. Phatnani, J. Shu, F. Parisi, E. Schmidt, P. Robins, L. Niederhofer, J. Wang, C. Aliferis

## Abstract

The work reported here establishes - for the first time in the literature - the existence of equivalence classes of Cellular Senescence (CS) signatures (and by extension of biomarkers and senotypes relating to those signatures) across various phenotypes, tissues, assay types and research designs. This means that typically there is no single CS signature (for a phenotype of interest) but many equivalent ones. This is true for both optimal (maximally predictive and non-redundant signatures) and non-optimal signatures. A consequence of this finding is the need for discovery of all equivalent signatures and making biological sense of them in toto. A variety of useful algorithmic methods exist to do so. Good modeling and research practices (such as ones proposed here) must accommodate the existence of CS equivalent signatures and associated biomarkers and senotypes.

## 1. INTRODUCTION

Cellular Senescence (CS), the phenomenon whereby cells prematurely and permanently stop dividing due to aging, stress, or damage), is a major hallmark of aging, and is linked to aging-related disease and loss of function. In recent years significant progress has taken place in mapping of CS across various tissues. The field is rapidly advancing from *Description* (where CS occurs) to *Classification and Prediction* (linking CS markers to cell, tissue, organ, organism phenotypes), toward *Causal (mechanistic) understanding* (what triggers CS and what are its consequences^1^. Machine Learning (ML), statistics, and other forms of complex modeling are essential tools for such discovery^2,3^.

Applications of *Predictive modeling (PM) in CS* for example include exploring heterogeneity of CS^4^; predicting the onset of mobility disability^5^; detection of CS cells^6^; Anti-senescent drug screening^7^; prognosis and therapeutic response of hepatocellular carcinoma^8^; using the senescence-associated secretome as an indicator of age and medical risk^9^; detecting the dynamic states of cells^10^; predicting their fates^11^, etc.

CS Predictive modeling includes *Classifiers and Regressors*. For example, using single cell transcriptomic measurements, a classifier may determine if a cell is senescent or not (or the probability that it is). A **regressor** predicts continuous instead of categorical responses (e.g., expected longevity on the basis of a CS or other “biological clock”^12^).

The concepts of *signatures, gene lists, molecular profiles, markers, biomarkers, features, feature selection*, etc, are often confused. In the present work marker sets of biological interest are referred to *as signatures*. A gene list is a common type of signature, however a signature may contain other (and often several) types of molecular species. An important distinction is between literature-derived vs data-driven signatures.

*Feature selection* (aka variable or marker selection, in the context of CS and aging research), methods enable us to discover signatures from data. This modeling stage is central to all high dimensional data modeling. It aims to reduce a large number of measured variables (from thousands to up to millions) down to small sets with ideally all the predictive information about the response variable. This makes predictive and causal (and other types of) modeling feasible/easier and may also reveal important markers for CS^13–17^.

*Example CS signatures include* SenMayo^18^ which is a signature that aims to identify senescent cells across tissues and species with high fidelity. Using this senescence panel, the authors were able to characterize senescent cells at the single cell level and identify key intercellular signaling pathways. Sensig^19^ is an in vivo-derived senescence signature using a foreign body response-driven fibrosis model in a p16-CreERT2;Ai14 reporter mouse. As a third example, a signature comprising 68 differentially expressed RNAs that helps to discriminate between senescent and non-senescent cells was reported in Casella, 2019^20^ (also discussed in Gorgoulis, 2019^21^). In Wechter, 2023^22^, signatures are used as part of the effort to identify different senescence programs that developed divergently, one in which cells expressed traditional senescence markers such as p16 (CDKN2A) mRNA, and another in which cells expressed long noncoding RNAs and splicing was dysregulated and to obtain evidence that the proliferation status at the time of senescence initiation affects the path of senescence, as determined based on the expressed RNAs. Finally the review of Li, 2025^23^ discusses use of signatures to detect organ- and cell type-specific senescence, since senescent cells exhibit marked heterogeneity across tissues due to differences in developmental origin, cell composition, extracellular milieu and exposure to age- or stress-induced damage. Investigating cellular senescence signatures, helps in the understanding of organ-specific contexts due to the diverse biological functions and cellular compositions of different tissues.

Particularly important for the present research is the relatively newly discovered phenomenon of signature information equivalence, predominantly established/studied in cancer research^24–28^, but mathematically possible in all omics data.

### Problems caused by signatures equivalence classes

In practical terms, having signatures/markers panels with non uniqueness^24,29^ (i.e., equivalence class >1 member) creates a number of significant problems. These problems are “invisible” to researchers who do not extract the signatures’ equivalence classes. Such problems include:

(a*) CS specificity*: Some signatures maybe CS-specific while other that are information equivalent may not be CS-specific. (b*) Causal interpretation* even when using causal algorithms with guarantees for reliable causal discovery is problematic because the TIE phenomenon violates the common fundamental algorithmic assumptions. (c) Similarly, the *estimation of causal effects of true causal factors* of phenotypes of interest becomes more complicated because they may become statistically indistinguishable from IE non-causal factors. (d) Large equivalence classes can also *confuse the meaning of statistically stable/unstable results, and tests of signature randomness*. (e) *Priority claims and IP protection* can be bypassed because equivalent signatures are essentially (predictively) “bioequivalent” diagnostics and theranostics. (f) Non-uniqueness of optimal signatures also creates the possibility of *apparent but false disagreement/non reproducibility of results among different groups*, when in reality they may be discovering fully equivalent signatures. (g) Finally the existence of large equivalence classes of signatures for the same phenotype *“breaks down” connectivity map* approaches that seek to connect candidate drugs such as small molecules and genetic reagents, to diseases using signatures as the query against a massive reference database of cellular responses induced by the drug candidates.

### Hypotheses

In the present research, and focusing on CS research, we investigate two interrelated hypotheses:

- *Hypothesis 1 (existence of predictive signal for phenotypes of interest):* CS signatures both literature-driven and data-driven contain information (measured e.g., as predictive signal) for important clinical and experimental phenotypes.
- *Hypothesis 2 (equivalent predictive signals):* CS signatures are not unique, i.e. they have equivalence classes with >1 (possibly numerous) members.

In the present work we first provide strong evidence that CS signatures (both literature and data driven ones) have strong signals for a range of studied phenotypes. We establish that literature-driven signatures can have extensive redundancies and in some cases smaller signals compared to data-driven ones. We also establish, for the first time in the literature, that both kinds of CS signatures are not unique and as a matter of fact they exhibit surprisingly large equivalence classes. We finally show that the literature-derived signatures in the majority of cases fail to pass statistical tests comparing them to random signatures of similar size, despite them being reproducible - as demonstrated by cross and holdout validation.

On one hand, the evidence in favor of Hypothesis1 strengthens the belief that CS can be a translational and clinical phenomenon with extensive impact for aging and Geroscience . On the other hand, the evidence in favor of Hypothesis 2, introduces serious complications regarding the CS specificity of markers/signatures and their mechanistic interpretation (e.g., toward drug target development or developing other deep biological understanding), along with the many problems of signature non-uniqueness enumerated above..

In the remainder we discuss our methods, data and experimental findings and propose corresponding good practices for navigating the problem of non-uniqueness of CS signatures.

## 2. METHODS

### 2.1. Operational definitions

We define a biomarker as and a signature as any molecular entity in omic data that is studied in the context of its predictive or causal relationship to phenotypes (e.g., biological function, disease or other) of interest. The term “biomarker” is within our scope but has a narrower interpretation since it typically relates to proxies of outcomes or indicators of side effects of therapeutic interventions^30,31^. An optimal predictive signature is considered one that has max predictivity and no redundancy (aka “minimality”). In addition, in several research contexts we require that optimal signatures also have causal (i.e. mechanistic) interpretability.

We defined signature TIE (target information equivalence) or equivalence the existence of more than one optimal signature in some distribution and/or the data sampled from it^29^. The existence of signature equivalence classes is revealed using specialized algorithms.

*Senotypes*^*32,33*^. *T*he senotype classification framework was developed by researchers in the NIH SenNet Consortium, recognizing that senescent cells are highly diverse depending on the tissue, the type of stress that caused them to stop growing, and their age. **The senotype classification uses as key dimensions:** Cell identity and context, inducing mechanism, temporal stage, multimodal molecular and structural features. *It is evident that insofar signatures and senotypes use common features, there is a correspondence between signatures and senotypes defined by their intersecting characteristics*.

A recent set of proposed good practices for developing validating and clinically deploying signatures is presented in Simon and Aliferis^34^.

**SUPPLEMENT A** provides technical details and examples: **A.1** signatures and signature optimality. **A.2**. Signature TIE equivalence. **A.3** algorithms for signature discovery, optimal signature discovery and equivalence classes discovery.

### 2.2. Predictive Modeling

The clinical phenotype in all tasks were dichotomous. We derived models to predict these phenotypes using bulk, aggregated single cell, and single cell data listed in Table 1. We divided each dataset into a Discovery set (70% of the samples) and Hold-out Validation set (30% of the samples). We used the Discovery set to build predictive models with the stratified repeated nested cross validation (NCV) protocol to ensure robust model selection and unbiased performance estimation. We conducted model selection in the inner loop of the NCV, and performance estimation in the outer loop of the NCV. Subsequently and independently, the model performance was validated on the Hold-out Validation set. We examined three model selection schemes for deriving de novo signatures, optimizing for predictive performance and parsimony with different levels of trade offs. The above protocol is statistically unbiased (i.e., exhibit no overfitting of error estimates); they allow for extensive model selection (hence reducing under-fitting). The “stacking” of nested cross-validation and hold-out validation with regularization of the classifiers employed and strong feature selector algorithms provided four layers of protection against overfitting^34^.

**Table 1.** Data used.

| Dataset | Outcome | # Samples | % Positives | # Genes | Data Type |
|---|---|---|---|---|---|
| GSE33000<sup>36</sup> | Alzheimer's | 467 | 66.38 | 39280 | Bulk |
| MITROSMAP<sup>37</sup>_AST | Alzheimer's | 306 | 52.29 | 24503 | Aggregate Single Cell |
| MITROSMAP_IMM | Alzheimer's | 306 | 52.29 | 21074 | Aggregate Single Cell |
| Parabiosis<sup>38</sup>_Kidney | Determine if organ-specific cells originate from attached (parabiosis) aged-aged mice pair (IA) vs young-young mice pair (IY) | 772 | 36.53 | 54444 | Single Cell |
| Parabiosis_Liver |  | 1013 | 19.35 | 54444 | Single Cell |
| Parabiosis_Thymusl |  | 1156 | 39.27 | 54444 | Single Cell |
| Parabiosis_Kidney | Determine if organ-specific cells originate from aged mice in aged-aged (IA) mice pair vs. aged-young mice pair (HA). | 504 | 55.95 | 54444 | Single Cell |
| Parabiosis_Liver |  | 504 | 38.89 | 54444 | Single Cell |
| Parabiosis_Thymus |  | 806 | 56.33 | 54444 | Single Cell |

### 2.3. Equivalence Class discovery

All equivalence class discovery methods employed in the current study involve the following two steps: (1) Derive the first irreducible feature set: Given an input feature set, derive one subset of features that is predictively optimal with respect to the input feature set and is irreducible (i.e. removing features from the subset results in decrease in predictive performance). For the literature derived signatures, the input feature set is the signature. For the de novo feature set, the input feature set is all the features in the dataset. (2) Derive additional predictively equivalent feature sets: Given the first irreducible feature set, identify predictively equivalent feature sets from all the features in the datasets. Different equivalence class methods/variants make different assumptions about the equivalence structure and utilize different hyper parameters. The equivalence class was derived using the discovery set and their performances validated in the validation set. See Supplement A for details.

### 2.4. Functional annotation

We compared our de novo derived signatures from equivalence class discovery to a list of 866 CS genes reported in prior literature which are archived and manually curated in CellAge, the database of cell senescence genes^35^.

### 2.5. Data

We examined 9 dataset-phenotype combinations from 3 prior studies. The 3 studies measure molecular features using bulk, aggregated single cell, and single cell gene expression, respectively. 2 studies (3 prediction tasks/phenotypes) collect human data and 1 study (6 predictive tasks/phenotypes) are from mouse experiments. Details in Table 1.

### 2.6 Literature-Derived Signatures

**Table 2** lists the three literature-derived CS signatures examined in the current study.

**Table 2.** Literature-Derived Signatures.

| Signature | # Genes |
|---|---|
| SenUMN | 33 |
| SenMayo<sup>18</sup> | 125 |
| SenMayo mouse<sup>18</sup> | 118 |

## 3. RESULTS

### 3.1. Predictive signal of literature-derived and de novo signatures for the 9 dataset-phenotype combinations

Results in this section are illustrated in **Figure 1, with more details in Table 1** in **Supplement B, Column H and O**. In the human data, predictive performances of both literature-derived signatures and de novo ones, were uniformly much higher in the human bulk data than the human single cell aggregate data (on limited cell types). Predictivity estimated by the NCV in the former ranged from 0.94 to 0.97 AUC (average 0.96), denoting almost perfect diagnosis. Predictivity in the latter ranged from 0.60 to 0.73 AUC (average 0.65). Ceteris paribus, this indicates that the chosen cell types in MITROSMAP do not carry all the information contained in the bulk data. However, these data-phenotypes are useful to study equivalence classes in moderate to low signal settings as we will see.

**Figure 1.**
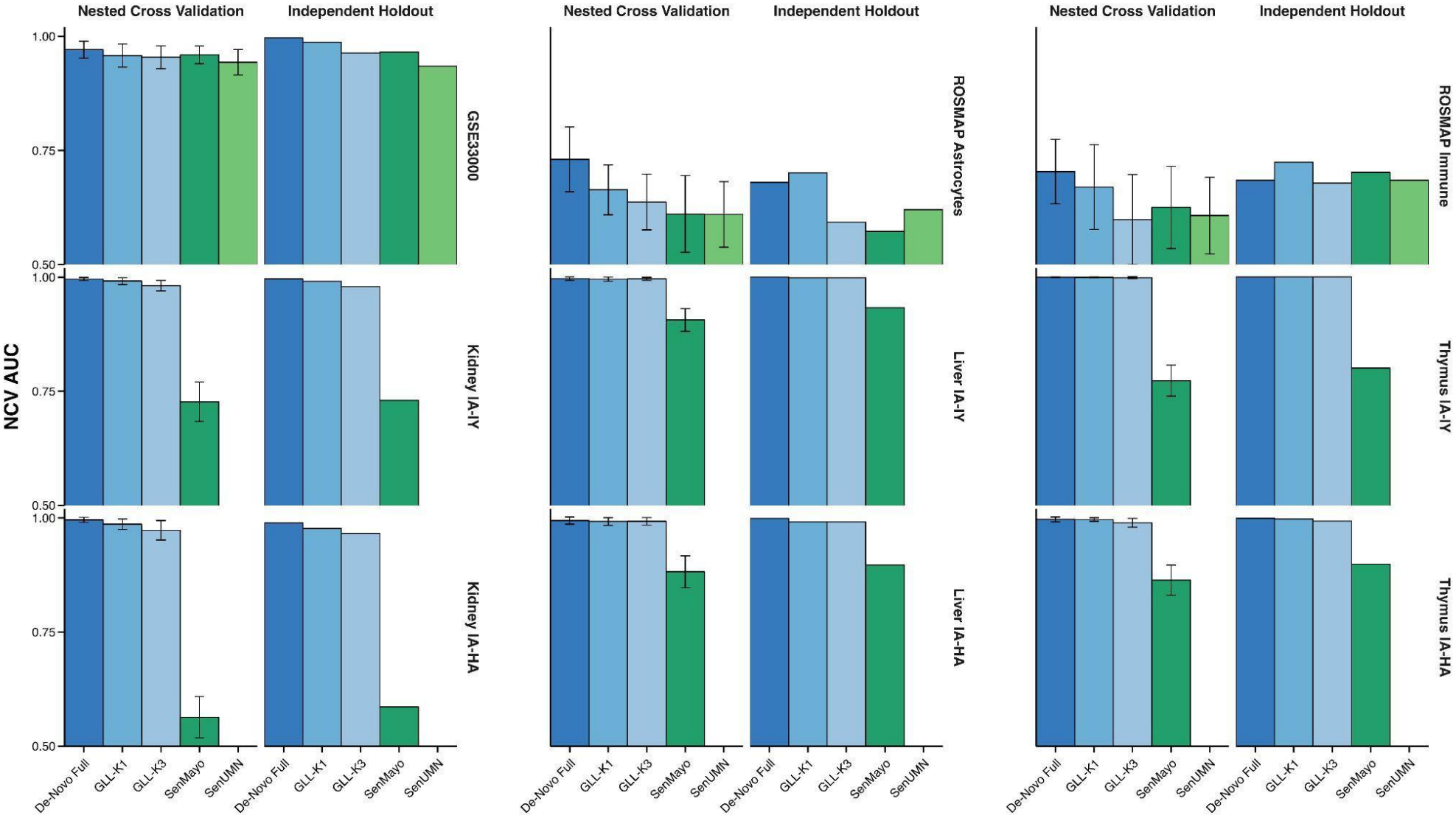
depicts both nested cross-validation predictivity estimates as well as additional independent validation estimates.

In the mice data, predictive performances were very high for de novo signatures, but less so for literature-derived ones. The former ranged from 0.97 to 1.00 (average 0.99). Predictivity in the latter ranged from 0.56 to 0.90 AUC (average 0.79).

In general, across all setups the literature derived signatures demonstrated mildly to strongly inferior predictive performances compared to de novo derived signatures.

### 3.2. Equivalent signatures, their predictivity, redundancy and randomness across the 9 dataset-phenotype combinations

We derived predictive equivalents for each dataset-phenotype combination for both the literature-derived signatures and for the de novo derived signatures.

We verified that the derived equivalent signatures perform the same as follows: we compared the estimated predictivities of single optimal data-derived signatures in the discovery datasets (as described in **Figure 2, and Table 2 Supplement B**, using nested cross validation) with the hold-out validation performance of each member of the corresponding equivalence class, see **Table 2 Supplement B Column M and P**).

**Figure 2.**
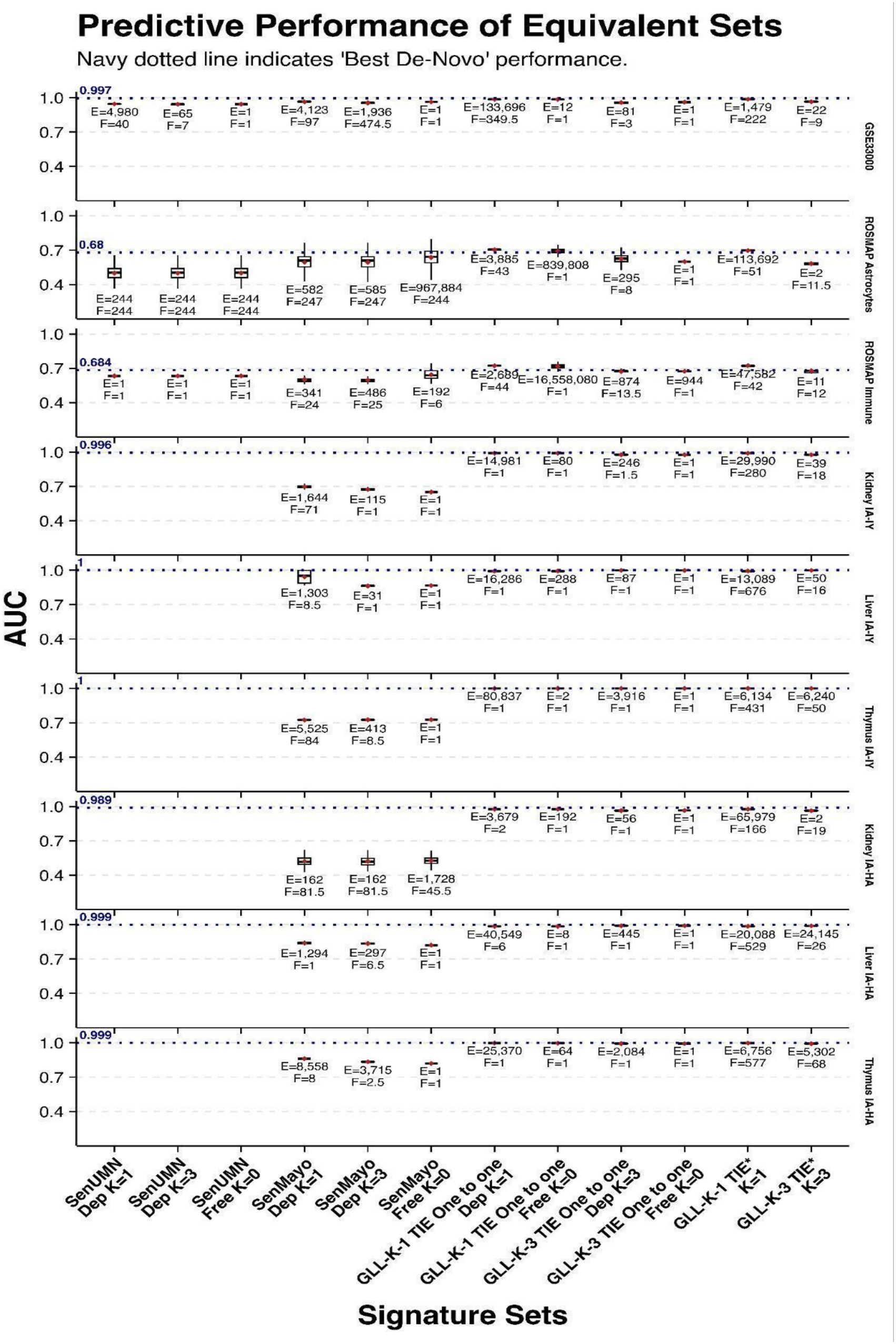
Predictive performance of equivalent sets.

Out of 90 equivalence class method-dataset-phenotype combinations, 72 (80%) exhibited non unique signatures. The rest 18 (20%) had unique signatures. The size of the 74 equivalence classes ranged from 2 to 16,558,080 members. See **Table 2 Supplement B column I**). We note that because our protocol eliminates redundancies from seed literature signatures before equivalents are induced from data, *the sizes of the equivalence classes can be very conservative*.

The literature-derived signatures exhibited *redundancy* for specific phenotypes. This redundancy is expected because they are not optimized for specific phenotypes (other than presences of CS). The redundancy was estimated by comparison to the most parsimonious equally predictive subset of the literature-derived signature in each data-phenotype combination and ranged from 75% to 98% (0% denoting no redundancy and 100% denoting all features redundant). See **Table 3 Supplement B Column K**.

As expected by their design, different equivalence class methods resulted in different numbers of equivalence signatures, because they trade off differently completeness (or data distribution applicability) with computational speed.

Finally, literature-derived signatures often ( 66% of the time, 8 out of 12 signature task combinations) were not statistically significantly different from random signatures of the same size. This is directly linked to their redundancy and the presence of predictively equivalent signatures. As shown in Ma,2026^39^, the higher number of predictively equivalent signatures (given fixed equivalence structure), the more likely randomly selected features will result in equivalent predictive performance as the original signature.See **Table 1 Supplement B Column S**.

### 3.3. Preliminary examination of biological functions of genes in the equivalence classes

Among all predictively optimal equivalent signatures, only one corresponds exactly to the local causal neighborhood (the rest may contain genes that are of the local causal neighborhood or no member at all; genes not on the local causal neighborhood may still be causal remotely, or they can be confounded)^29^. Under the assumption of *causal sufficiency* (i.e., no unmeasured confounders)^40^, if a gene appears in *all* equivalent signatures, it is guaranteed to be part of the local causal neighborhood^29,41^, however. Such genes become a natural first priority when interpreting results of data analysis.

Supplemental **Table 4 in Supplement B** lists all genes that appear in all equivalent signatures for at least one equivalence class discovery method, for some dataset-outcome combination. We designate those as “*priority causal relevant genes*” following the logic described above.

Resolving the remaining causal uncertainty requires prior knowledge and/or experiments. Algorithms do exist that try to minimize the number of such experiments, and guarantee correct discovery of the causal structure underlying the biological system under study^41^.

We compared these genes to a comprehensive list of 866 genes known related to senescence previously described in the CellAge database^35^. As shown in **Table 3**, for any dataset-phenotype combination, only a small percentage of the de novo genes (ranging from 1.88% to 11.13%) deemed to be *priority causal relevant* are actually known to be related to senescence. Of the 866 genes, 696 (80.37%) of the genes did not overlap with the priority causal relevant genes.

**Table 3.** Occurrences of CS genes in the priority causal relevance group (listed in Table 4, SB)

| **Dataset** | **Outcome** | **# Genes** | **# Priority causal relevant genes** | **# of CS genes in the Priority causal relevance group** | **% of CS genes in the Priority causal relevance group** |
|---|---|---|---|---|---|
| GSE33000 | Alzheimer's | 39280 | 222 | 9 | 4.05 |
| MITROSMAP_AST | Alzheimer's | 24503 | 41 | 1 | 2.44 |
| MITROSMAP_IMM | Alzheimer's | 21074 | 27 | 2 | 7.41 |
| Parabiosis_Kidney | IA vs IY | 54444 | 277 | 12 | 4.33 |
| Parabiosis_Liver | IA vs IY | 54444 | 674 | 75 | 11.13 |
| Parabiosis_Thymus | IA vs IY | 54444 | 431 | 30 | 6.96 |
| Parabiosis_Kidney | IA vs HA | 54444 | 160 | 3 | 1.88 |
| Parabiosis_Liver | IA vs HA | 54444 | 527 | 41 | 7.78 |

## 4. DISCUSSION

The importance of CS and its various pathways and markers for longevity and longevity-related ones such as loss of function and age-related disease is well documented in prior work^42–44^. Building molecular signatures capable of predicting such phenotypes is an important approach for studying the role and impact of CS, and can also have clinical potential (diagnostic, risk-stratifying, etc.).

The degree by which literature-derived - “CS omnibus signatures” - are informative across a range of for phenotypes - *which were not used for the construction of the signature* - was found to be unexpectedly high in the present study (given their phenotypic non-specificity). We also found that in all studied cases phenotype-specific signatures created de novo from related datasets (a) would match or exceed literature-derived ones, and (b) did not always comprise exclusively known

CS-specific markers. As expected, omnibus literature signatures are to large extent redundant for specific phenotypes because by design they contain markers that are useful across a range of phenotypes and are not tailored to particular phenotypes.

The most important finding of the present work. that to the best of our knowledge *is established here for the first time in the literature*, is that the predictively optimal and non-reducible senescence signatures developed in the majority of studied data and phenotypes are not unique. Their equivalence classes in some cases have many members even reaching in the thousands or millions of signatures. CS signatures exhibit large equivalence classes across several phenotypes, data designs, and omics types. Preliminary functional annotation shows that the phenotype-optimized equivalence classes contained non CS genes and CS ones. A more detailed analysis of the equivalence class is warranted, however and will be undertaken in a forthcoming extension of the present work.

The common and commonly large equivalence signature classes create several immediate problems: for the interpretation of signatures, for the synthesis of results across research groups, for the discovery of drug targets, for the in silico estimation of causal effects for a variety of markers, and for priority claims and protection of IP. It is, moreover, not clear at this time how much these equivalence classes have hindered or slowed down progress in the field of CS and longevity therapeutics, more broadly. Stated differently, being oblivious to equivalence classes of CS signatures introduces both False Positive and False Negative results, and it is not clear how many such false positive and false negatives occurred in the published senescence literature.

The non-uniqueness of CS signatures *also implies non uniqueness of senotypes*. Because CS signatures inherently (logically, and mathematically) relate to senotypes, for the corresponding phenotypes, an equivalence class of signatures entails the existence of an equivalence class of senotypes. This is a mathematical necessity; in the reported work we did not attempt to empirically show equivalence classes of senotypes. This will be a future extension of the current work.

Our experiments utilized several algorithms for discovery of CS signature equivalents, with complementary properties and trade-offs. The algorithms used include previously published families (notably TIE* and iTIE*) but also novel enhancements that provide guarantees of soundness and completeness in more specialized (restricted) distributions, and also allow for discovery of equivalence classes of seed signatures (e.g., omnibus literature-derived CS ones) even when these are not optimal (from a combined predictivity and parsimony perspective).

While the TIE* algorithm family is enumerative and worst-case exponential in both time and space, both iTIE* and the novel algorithms are low-order polynomial (quadratic), in space and time, offering massive advantages in storage and interpretation of the discovered equivalence classes, at the expense of false negatives (in some distributions). In the small sample, the various algorithms have different susceptibility to statistical errors that occur because of small sample size variance. The precise operation and tradeoffs, as well as approaches to combine results from different algorithms will be addressed in forthcoming extension to the present work. We observe, however, that the equivalent signatures discovered in the set of experiments reported here are statistically robust as estimated by nested cross-validation and in addition generalize well to independent datasets (hence false positives are limited across algorithms).

An important point is that signature equivalency from a predictive perspective can be verified or rejected in a straightforward manner with application to datasets not used to derive the signatures and estimating predictivity there with unbiased estimators. From a causal perspective, however, experimental validation (along with literature and pathway database cross-referencing) is of paramount importance. Algorithms exist (e.g., the ODLP* algorithm family) to guide such experimentation such that the number of experiments needed to recover the true local causal pathway around a target variable of interest, is minimized (or greatly reduced). The ODLP* experimentation guiding algorithm guarantees that the number of single-variable experiments will never exceed the size of the union of the equivalent signatures of the target variable. For example if the union comprises 1 million signatures over 60 genes (identified from, e.g., 10,000 genes in the data), the maximum number of single-gene experiments guided by the algorithm in order to learn the complex causal pathway (including all higher order interaction), will not exceed 60. Note that pursuing a purely experimental approach will require up to o(2^10,000^ ) joint experimental gene manipulations, rendering it utterly infeasible. The algorithm-driven experimental approach opens the road for efficient discovery of causal CS pathways even in the presence of large equivalence classes.

The work discussed here is the first part of our investigation in CS signature/biomarker and senotype bioequivalence, to be followed with a forthcoming manuscript delving into detailed biological interpretation of our findings and analysis of the results obtained by different algorithms.

Our findings support the *good modeling practices* addressing the above problems summarized in **Table 4**. .

**Table 4.**
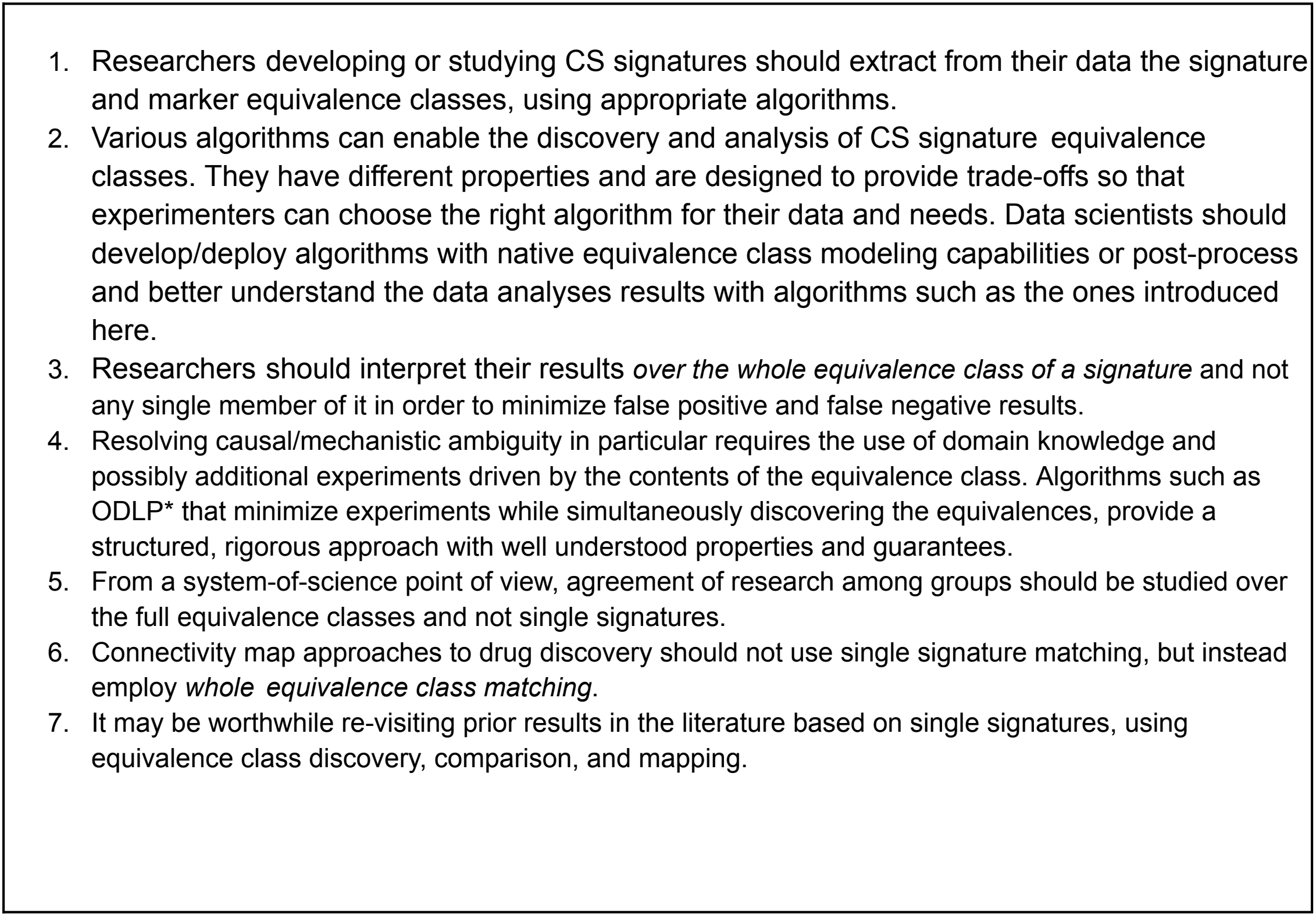
Proposed Good Modeling Practices based on the findings of the present study.

|  |
|---|
| 1. Researchers developing or studying CS signatures should extract from their data the signature and marker equivalence classes, using appropriate algorithms. |
| 2. Various algorithms can enable the discovery and analysis of CS signature equivalence classes. They have different properties and are designed to provide trade-offs so that experimenters can choose the right algorithm for their data and needs. Data scientists should develop/deploy algorithms with native equivalence class modeling capabilities or post-process and better understand the data analyses results with algorithms such as the ones introduced here. |
| 3. Researchers should interpret their results *over the whole equivalence class of a signature* and not any single member of it in order to minimize false positive and false negative results. |
| 4. Resolving causal/mechanistic ambiguity in particular requires the use of domain knowledge and possibly additional experiments driven by the contents of the equivalence class. Algorithms such as ODLP* that minimize experiments while simultaneously discovering the equivalences, provide a structured, rigorous approach with well understood properties and guarantees. |
| 5. From a system-of-science point of view, agreement of research among groups should be studied over the full equivalence classes and not single signatures. |
| 6. Connectivity map approaches to drug discovery should not use single signature matching, but instead employ *whole equivalence class matching*. |
| 7. It may be worthwhile re-visiting prior results in the literature based on single signatures, using equivalence class discovery, comparison, and mapping. |

**In conclusion**, the phenomena described here suggest that study of signatures and marker panels in the field of CS (as well as other omics and biomedical domains exhibiting equivalence classes of optimal signatures) in order to reduce errors and improve the quality and speed of their investigations, *should migrate from a single signature paradigm toward an equivalence class view of CS markers, signatures and related senotypes*.

## ACKNOWLEDGMENTS

We thank the following groups, people, and entities for supporting this work: the SenNet Working group on Good Practices: Analytical Methods and Modeling (GP-AMM); the Clinical and Translational Science Institute (BLS Histology); the University of Minnesota Genomics Center; the resources and staff at the University of Minnesota University Imaging Centers (UIC) SCR_020997. The authors acknowledge the Minnesota Supercomputing Institute (MSI) at the University of Minnesota for providing resources that contributed to the research results reported within this paper.

Professors Vesna Garovic, Marissa Schafer, Sandeep Khosla, Scott Budinger, and Peter Adams provided valuable initial advice for the work reported here, both as members of SnNet Steering Committee, and as members of the expert biologist collaborator group formed to analyze the biological implications of the findings presented here. That work will appear in a forthcoming manuscript addressing biological Interpretation and implications of CS signature equivalence classes for senescence research.

