## Supplement A for "Cellular Senescence Signatures Accurately Predict Clinical Phenotypes But Lack Uniqueness. Part I: Demonstrating Equivalence Classes of Senescence"

November 13, 2026

#### 1 Signatures and Signature Optimality

In the present report, biological marker or marker denotes any molecular entity in omic data that is studied in the context of its predictive or causal relationship to phenotypes (e.g. biological function, disease, or other) of interest. The term "biomarker" is within our scope but has a narrower interpretation, since it typically relates to proxies of outcomes or indicators of side effects of therapeutic interventions [1, 2]. A signature is a set of marker, but also implies the corresponding mathematical or computational model(s) that takes as input the markers' value and outputs the most accurate predictions that the data allows with regards to the phenotype of interest. Such models can be developed with standard model fitting procedures using classifiers and modeling protocols with empirical and theoretical optimal performance. The marker sets (sets of variables) that can result in models with optimal predictive performance for a phenotype of interest are Markov Blankets. A minimal (i.e. removing any variable will result in loss of information for the phenotype) Markov Blanket is a Markov Boundary.

**Definition 1: Markov Boundary of $T$ [3].** *A Markov boundary of $T$, denoted as $\mathbf{MB}(T)$, is a minimal set of features with the following property: $\forall \mathbf{S} \subseteq \mathbf{V} \backslash \mathbf{MB}(T)$, $\mathbf{S} \perp T | \mathbf{MB}(T)$.*

**Definition 2: Markov Blanket [3].** *A variable set $\mathbf{MB}(T)$ where $\mathbf{MB}(T) \subseteq \mathbf{MBL}(T)$ is a Markov blanket.*

Markov Boundaries and Markov Blankets are always anchored on a specific outcome or phenotype of interest. However, for brevity of notation, they are often denoted as **MB** and **MBL**, without specifically referencing the outcome variable.

Under the conditions that the learner that constructs the classification model can learn the distribution $p(T|\mathbf{MB})$, and that the loss function is such that perfect estimation of the probability distribution of $T$ is possible. Models built with **MB** results in optimal performance for $t$ [4].

### 2 Signature Target Information Equivalence

In faithful distributions there is one unique Markov Boundary for $T$, consisting of the parents, children, and parents of children of $T$ [5]. However, in distributions with faithfulness violations, there can exist multiple Markov Boundaries, leading to multiple optimal models/signatures [6].

**Definition 3: Markov Boundary Conditional Target Information Equivalence (MBC-TIE).** *Feature sets* $\mathbf{S}_1$ *and* $\mathbf{S}_2$ *are Markov boundary conditional target information equivalent when there exist two Markov boundaries of* $T$*,* $\mathbf{MB}_1$ *and* $\mathbf{MB}_2$*, such that:* $\mathbf{S}_1 \subset \mathbf{MB}_1$*,* $\mathbf{S}_2 \subset \mathbf{MB}_2$*,* $\mathbf{MB}_1 = \mathbf{S}_1 + \mathbf{MB}_1 \cap \mathbf{MB}_2$*,* $\mathbf{MB}_2 = \mathbf{S}_2 + \mathbf{MB}_1 \cap \mathbf{MB}_2$*. These equations are referred to as substitution functions, since they specify that one can obtain*$\mathbf{MB}_2$ *by substituting* $\mathbf{S}_1$ *with* $\mathbf{S}_2$ *in* $\mathbf{MB}_1$ *and vice versa.*

### 3 Algorithms and Analysis Pipeline for Signature Discovery, Optimal Signature Discovery, and Equivalence Class Discovery

#### 3.1 Algorithms and Analysis Pipeline for Signature Discovery

In the current study, the clinical phenotype for all tasks were dichotomous. We derived models to predict these phenotypes using bulk, aggregated single cell, and single cell data (Table 1 in the manuscript). We divided each dataset into a Discovery set (70% of the samples) and Hold-out Validation set (30% of the samples). We used the Discovery set to build predictive models with the stratified repeated nested cross validation (NCV) protocol to ensure robust model selection and unbiased performance estimation. We conducted model selection in the inner loop of the NCV, and performance estimation in the outer loop of the NCV. Subsequently and independently, the model performance was validated on the Hold-out Validation set. We examined three model selection schemes for deriving de novo signatures, optimizing for predictive performance and parsimony with different levels of trade offs by allowing the selection among different subsets of feature selection methods. The three model selection schemes are the following: (1) selection among all feature selection methods (see a list of all feature selection methods below), (2) GLL-PC feature selection with K=1, (3) GLL-PC feature selection with K=3. These three model selection schemes places progressive emphasis on model parsimony and results in models with progressively smaller number of input variables.

The above protocol is statistically unbiased (i.e., exhibit no overfitting of error estimates); they allow for extensive model selection (hence reducing underfitting). The "stacking" of nested cross-validation and hold-out validation with regularization of the classifiers employed and strong feature selector algorithms provided four layers of protection against overfitting [7].

For de-novo signature discovery, we used four six selection methods: all features, fisher, support vector machine recursive feature elimination, Rnadom forest recursive feature elimination, lasso, and GLL (generalized local learning, which infers MBs of the outcome) [5], and four classifiers: logistic regression, random forests, Boosted Trees, and support vector machines, chosen because of their theoretical properties and excellent empirical performance across many biomedical domains/datasets and especially clinical and omics biomolecular data in relatively low sample size domains [5, 7]. The hyper-parameter for the GLL was K= [1, 3]. The hyper-parameter for random forest was 500 or 1000 trees, mtry=sqrt (number of variables). The hyper-parameter for boosted tree was 500 or 1000 trees, learning rate of 0.1, using a tree stump. For support vector machines, we used the Cartesian product of the following hyper parameters: polynomial kernel degree= [1,2,3] and C=[0.1,1,10]. Selection among models built with the above combinations of feature selection and classication methods were conducted in the innerloop of the NCV. Missing data were handled by mean imputation, and we included indicator variables for missingness. The imputation was done according to the training data at all stages of the model development and performance estimation to prevent information leakage between train-test splits leading to overfitting.

##### 3.2 Optimal Signature Discovery

As discussed above, in theory, the Markov Boundary of a target variable of interest results in predictively optimal models with the minimal set of features. Therefore, our primary method for optimal signature discovery is to derive Markov boundaries using GLL algorithm variants [5].

##### 3.3 Equivalence Class Discovery

All equivalence class discovery methods employed in the current study involve the following two steps: (1) Derive the first irreducible feature set: Given an input feature set, derive one subset of features that is predictively optimal with respect to the input feature set and is irreducible (i.e. removing features from the subset results in decrease in predictive performance). For the literature derived signatures, the input feature set is the signature. For the de novo feature set, the input feature set is all the features in the dataset. (2) Derive additional predictively equivalent feature sets: Given the first irreducible feature set, identify predictively equivalent feature sets from all the features in the datasets. Different equivalence class methods/variants make different assumptions about the equivalence structure and utilize different hyper parameters.

We used a variety of methods from the TIE* (discovers Markov boundaries with many-to-many equivalence structure), and iTIE* algorithm family (discovers Markov Boundaries with one-to-many equivalence structure) [6]. In Figure 2 of the manuscript, methods with names containing "TIE*" belong to the TIE* algorithm family, methods with names containing "one to one" belongs to the iTIE* algorithm family. Some variants of the iTIE* family was deployed in the current study for the first time, enabling the identification of equivalence classes for non-optimal feature sets. We used these variants of the algorithm to identify equivalence classes for the literature derived signatures. Detailed description of these iTIE* variants will be described in a future technical paper.

The equivalence classes was derived using the discovery data and their performances validated in the validation set.

[3] Judea Pearl. *Probabilistic Reasoning in Intelligent Systems: Networks of Plausible Inference*. San Francisco, CA, USA: Morgan Kaufmann Publishers Inc., 1988. ISBN: 1558604790.

[4] Ioannis Tsamardinos and Constantin F Aliferis. "Towards principled feature selection: Relevancy, filters and wrappers". In: *International Workshop on Artificial Intelligence and Statistics*. PMLR. 2003, pp. 300–307.

[5] Constantin F Aliferis et al. "Local causal and Markov blanket induction for causal discovery and feature selection for classification part I: algorithms and empirical evaluation." In: *Journal of Machine Learning Research* 11.1 (2010).

[6] Alexander Statnikov et al. "Algorithms for discovery of multiple Markov boundaries". In: *Journal of Machine Learning Research* 14.Feb (2013), pp. 499–566.

[7] Gyorgy J Simon and Constantin Aliferis. "Artificial intelligence and machine learning in health care and medical sciences: best practices and pitfalls". In: (2024).
